# PIM_6_ Induces Tolerance and Confers Protection against *Mycobacterium tuberculosis* in Primary Macrophages

**DOI:** 10.64898/2026.11.03.763869

**Authors:** Maia Serene Gower, Linn Kleberg, Carolina Sousa Silva, Martine Gilleron, Jérôme Nigou, Margarida Correia-Neves, Christopher Sundling, Gunilla Källenius

**Affiliations:** Division of Infectious Diseases, Department of Medicine Solna, Karolinska Institutet, Stockholm, Sweden; Department of Infectious Diseases, Karolinska University Hospital, Stockholm, Sweden; Center for Molecular Medicine, Karolinska University Hospital, Stockholm, Sweden; Univ Toulouse, CNRS, IPBS, Toulouse, France; Life and Health Sciences Research Institute (ICVS), School of Medicine, University of Minho, Braga, Portugal

**Author notes:** **Corresponding author:** Christopher Sundling.

## Abstract

Glycolipids are major components of the mycobacterial cell envelope and play critical roles in host-pathogen interactions. Here we investigate the capacity of the mycobacterial glycolipid phosphatidyl-myo-inositol hexamannosides (PIM_6_) to modulate innate responsiveness in naive primary human monocytes and monocyte-derived macrophages (MDMs). Monocytes and MDMs prestimulated with PIM6, followed by 5 days of rest, secreted attenuated levels of tumor necrosis factor alpha (TNF-α), interleukin 6 (IL-6), and interleukin 10 (IL-10) following LPS challenge. PIM_6_ pretreatment significantly altered pattern recognition receptor expression through the experimental time course and programmed a tolerized phenotype in myeloid cells independent of TLR2 activity. In an infection challenge with H37Rv *Mycobacterium tuberculosis* (M.tb), MDMs pretreated with PIM_6_ secreted decreased levels of TNF-α and IL-10 and exhibited early increased bacterial control. Together, these data reveal PIM_6_ as a complex immunomodulator that tolerizes myeloid cells to subsequent stimuli and exerts direct effects on the control of M.tb infection.

## Introduction

The mycobacterial cell envelope is a complex and stratified structure unique to the mycobacterial genus. An inner plasma membrane composed of phospholipids and phosphatidylinositol-derived glycolipids including phosphatidyl-myo-inositol mannosides (PIMs), lipomannans (LM) and lipoarabinomannan (LAM), is encapsulated by a stratum of peptidoglycan covalently attached to arabinogalactan. The inner leaflet of the outer membrane is made of long-chain mycolic acids that esterify the terminal arabinose residues of arabinogalactan. The outer leaflet is then composed of a diverse array of non-covalently linked glycolipids including PIMs, LM, LAM, and other lipid families such as sulfoglycolipids, mycolic acids, trehalose-, and mycoserosate-containing lipids (Daffé & Marrakchi, 2019; Minnikin & Brennan, 2020). Due to their abundance and location in the outer layers of the mycobacterial cell envelope, these lipids are among the first molecules sensed by the innate immune cells, underpinning their importance in TB sensing and pathogenesis (Correia-Neves *et al*, 2022) .

PIMs and their biosynthetic derivatives LM and LAM share a common mannosyl-phosphatidylinositol anchor. The PIMs family comprises mono-to hexa-mannosylated forms (PIM_1 to 6_) but PIM_1_ and PIM_2_ are the most abundant glycoforms found in the mycobacterial cell envelope, including *Mycobacterium tuberculosis* (M.tb) (Daffé & Reyrat, 2008; Angala *et al*, 2014; Gilleron *et al*, 2003, 2014). PIMs and LAM have been shown to be distinct immunomodulatory entities in M.tb infection through blockade of phagosome maturation, downmodulation of IL-10, uncontrolled upregulation of proinflammatory cytokines, and type-1 interferons in monocytes, macrophages and dendritic cells in both *in vivo* and *in vitro* systems (Gilleron *et al*, 2014; Daffé & Reyrat, 2008; Vergne *et al*, 2015; Correia-Neves *et al*, 2022; Mishra *et al*, 2011).

We previously demonstrated that myeloid cells from Interferon gamma release assay (IGRA) positive individuals have an attenuated cytokine production *ex vivo* in response to M.tb glycolipids, when compared to cells from those with tuberculosis (TB) disease and healthy controls (Silva *et al*, 2021). From this observation, we hypothesized a mechanism in which a tolerized innate immune memory phenotype elicited by some component, such as the immunomodulatory glycolipids of M.tb, could function as a programmer of this dampened innate response.

In this study, we utilized stimulation protocols developed in the field of innate immune memory to explore the immunomodulatory effects of M.tb glycolipids on primary human myeloid cells. This consisted of initial stimulus with glycolipids, followed by a challenge with LPS 5 days after stimulus. This assay directly mimics other works evaluating innate training. Tumor necrosis factor alpha (TNF-α), interleukin-6 (IL-6), and interleokin-10 (IL-10), cytokines important in TB pathogenesis and dictation of both pro-and anti-inflammatory response in cells (Kireev *et al*, 2025; Roach *et al*, 1993; Flynn *et al*, 1995; Flynn & Chan, 2001; Cooper *et al*, 2011; Orme *et al*, 2015; Redford *et al*, 2011), were quantified upon this restimulation with LPS or PIM_6_. We further investigate if mycobacterial glycolipids alter the expression of pattern recognition receptors on myeloid cells over time and investigate TLR2 as a potential receptor for inducing the tolerized phenotype. Finally, we probe the functional effect of glycolipid exposure to control of subsequent M.tb infection in an in vitro infection model of glycolipid-pretreated monocyte-derived macrophages.

## Results

### PIM_6_-Stimulated Primary Myeloid Cells Downregulate Cytokine Production in Subsequent Stimuli

Following previous studies (Bekkering *et al*, 2016; Vuscan *et al*, 2024; Ifrim *et al*, 2014; Netea *et al*, 2020; Domínguez-Andrés *et al*, 2021), we established a protocol by which mycobacterial glycolipid-induced innate memory could be probed in primary human myeloid cells (**Figure 1A)**. Primary monocytes were isolated to a high level of purity to avoid T-cell contamination effects on myeloid receptor expression and response to stimuli (Nielsen *et al*, 2020; Elkord *et al*, 2005). Monocytes were isolated from freshly collected buffy coats from healthy blood donors using CD14^+^ magnetic bead selection, as this selection method yielded the least CD3^+^ contamination when compared to plastic adherence or CD14 negative magnetic selection via gating of CD33^+^CD14^+^ live cells (**Figure S1**). A threshold of 95% monocyte purity was used for subsequent stimulations. Monocytes were differentiated with GM-CSF (M1Ms), M-CSF (M2Ms), or GM-CSF+IL-6 (MDDCs) for six days prior to stimulation and verified for differentiation as in previous protocols(Mily *et al*, 2020).

**Figure 1.**
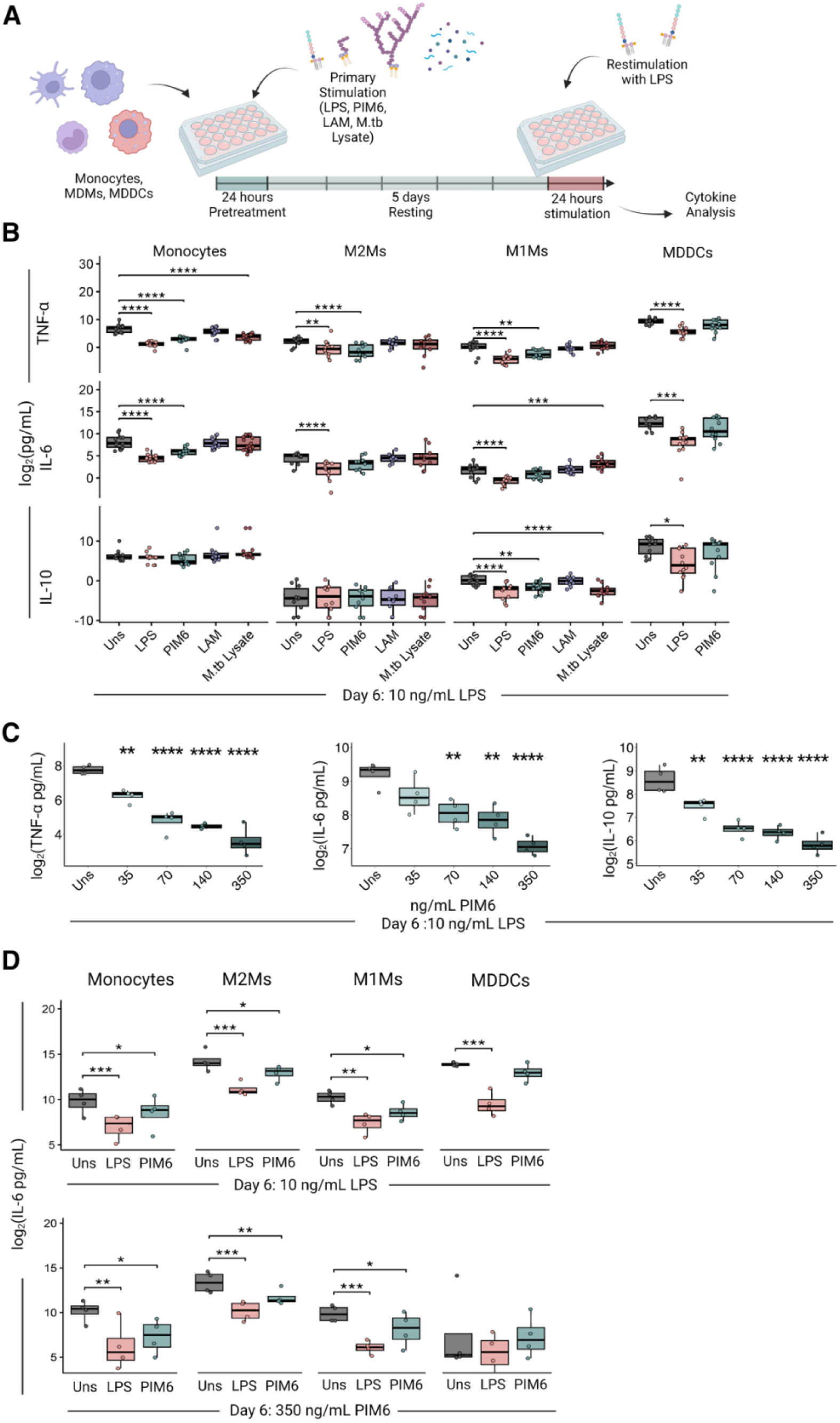
Cytokine secretion from LPS restimulation of glycolipid-pretreated myeloid cells indicates cell subset-specific and dose-dependent tolerogenic effects of PIM_6_. **(A)** Schematic illustration of the stimulation protocol to evaluate training and tolerance in primary monocytes, MDMs, and MDDCs. **(B)** Myeloid cells were stimulated for 24 hours with LPS, PIM_6_, LAM or M.tb Lysate before washing and resting for 5 days. Following day 6 LPS restimulation, TNF-α, IL-6, and IL-10 levels were quantified in the supernatant from all cell types. Post-hoc pairwise comparisons were conducted using a paired t-test with Bonferroni correction, (n=16). **(C)** Different concentrations of PIM_6_ were used for primary stimulation and cytokine analysis was conducted from following LPS stimulation, (n=4). **(D)** Monocytes were stimulated with differing concentrations of PIM_6_ (350 ng/mL, 140 ng/mL, 70 ng/mL, and 35 ng/mL) and TNF-α, IL-10, and IL-6 levels were quantified against unstimulated control upon day 6 LPS or PIM_6_ stimulation, (n=4). Statistics: One-way repeated measures ANOVA. Ns: p > 0.05, *: p ≤ 0.05, **: p ≤ 0.01, ***: p ≤ 0.001.

Monocytes and monocyte-differentiated cells were stimulated for 24 hours with LPS, a PIM_6_ enriched glycolipid fraction, LAM-enriched purified glycolipid fractions (LAM), or M.tb whole cell lysate or left unstimulated in complete media (Uns). After stimulation, cells were washed and rested for 5 days in complete RPMI with media changes every 48 hours. On Day 6, to assess cells’ ability to respond to subsequent challenge, cells from all conditions were stimulated with LPS for 24 hours and supernatant was collected for quantification of TNF-α, IL-6, and IL-10.

Monocytes and both subsets of MDMs stimulated with PIM_6_ showed a dampened cytokine production of TNF-α. While diminished secretion of IL-6 and IL-10 was observed in monocytes and M1Ms respectively, indicating a glycolipid-modulated immune response (**Figure 1B**). These results were recapitulated in primary human monocytes stimulated with PIM_6_ sourced from BEI resources or isolated in house using previously described methods (**Figure S2**)(Gilleron *et al*, 2003, 2001). Cell quantity and viability were consistent between conditions, as determined by Lactate-Dehydrogenase assay and ATP production assay either immediately prior to or following the day 6 LPS stimulation in monocytes (**Figure S2**).

In innate immune memory models, ligand concentration can determine either a trained or tolerized phenotype (Ifrim *et al*, 2014; Lajqi *et al*, 2023). To investigate if this feature applied to M.tb glycolipid stimulation, we stimulated monocytes with a series of PIM_6_ concentrations. Increasing concentrations of PIM_6_ led to progressively decreased secretion of TNF-α, IL-10 and IL-6 in response to day 6 LPS stimulation **(Figure 1C)**, indicating that PIM_6_-induced tolerization was dose-dependent.

LPS is a common ligand used to challenge if initial stimulus with an antigen results in induction of innate memory. Importantly, an innate training or tolerizing phenotype is not reliant upon the identity of the challenge (Netea *et al*, 2020; Saeed *et al*, 2014). To investigate if this tolerized restimulation phenotype was antigen agnostic, we conducted parallel experiments in which either LPS or PIM_6_ were used as stimuli on day 6 stimulation. Cytokine secretion analysis following this secondary stimulation demonstrated that PIM_6_ stimulated monocytes and MDMs demonstrated a dampened TNF-α and IL-6 secretion, regardless of rechallenging antigen (**Fig 1D**).

### PIM_6_ Induces Sustained Changes in PRR Expression in Primary Human Myeloid Cells

To assess whether stimulation with PIM_6_ alters the receptor expression profile of myeloid cells, we longitudinally characterized the expression of key pattern recognition receptors (PRRs) involved in recognition of mycobacterial glycolipids, alongside cell identity markers at distinct timepoints in our protocol. Expression of CD14, CD16, DC-SIGN, Dectin-2, Mincle, TLR2, TLR4, TREM2, and hDCIR was profiled with spectral flow cytometry upon seeding of monocytes or differentiation for MDMs and MDDCs, following primary stimulation with PIM_6_ or LPS (Stim.), as well as directly before (Resting), and after secondary stimulation with LPS on day 6 (Re-stim.). PIM_6_ and LPS stimulation led to modulated expression of several PRRs throughout our protocol, indicating stimulation-dependent effects on the cells (**Figure 2A, B, S3**). For PIM_6_ treated cells, TLR2 expression was consistently elevated in monocytes and M2Ms (**Figure 2B-D**). Dectin-2, DC-SIGN, and Mincle also showed altered expression over time.

**Figure 2.**
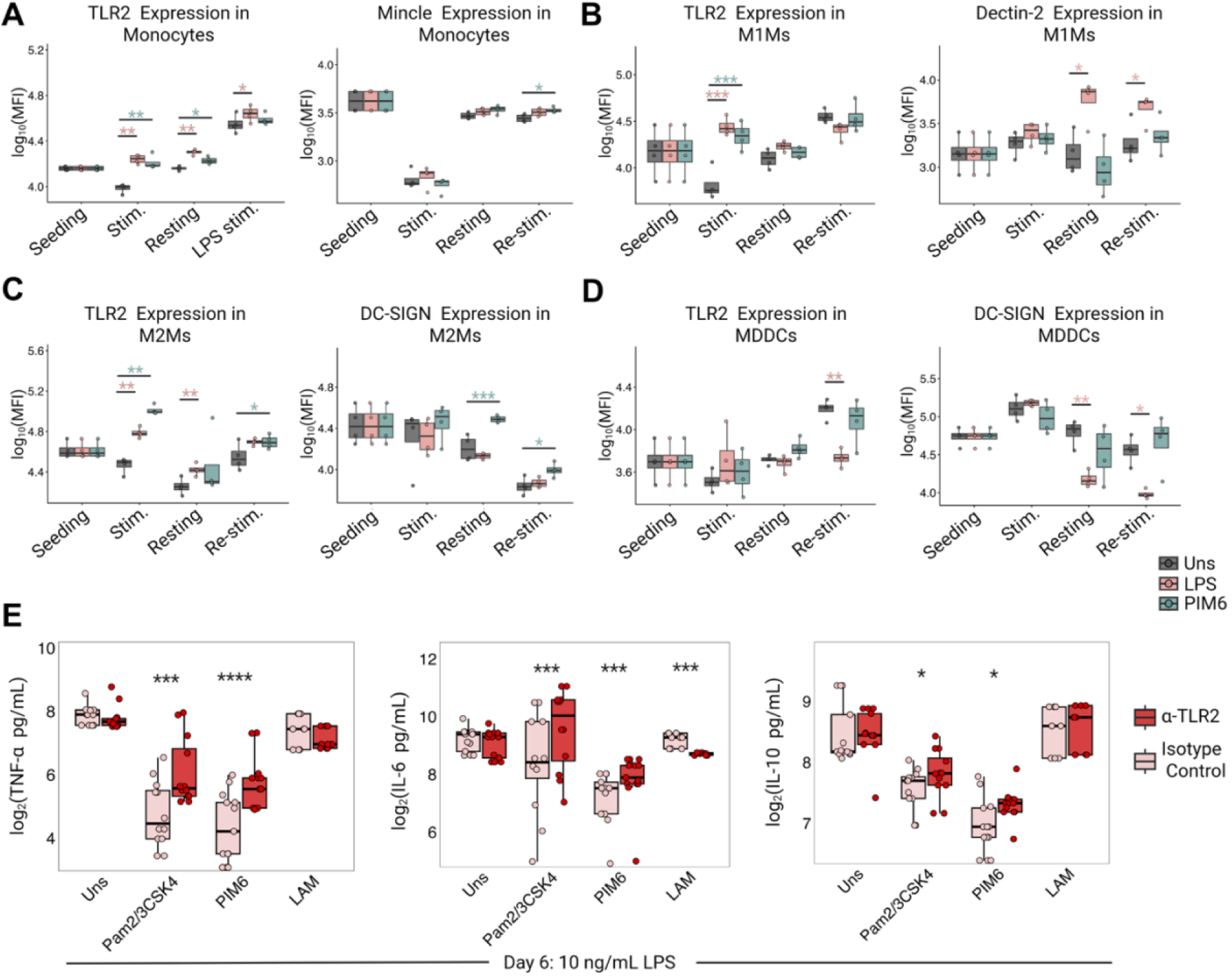
Pattern recognition receptor expression on primary myeloid cells is modulated in response to glycolipid stimulus. (**A-D**) Expression (MFI) of pattern recognition receptors were analyzed via flow cytometry at cell seeding, following glycolipid and LPS stimulus (Stim.), and immediately before (Resting) and after LPS restimulation (Re-stim.) (n=4). (**A**) MFI of Mincle and TLR2 expression in monocytes through course of experiment. (**B**) MFI of TLR2 and Dectin-2 in M1Ms through course of experiment. (**C**) MFI values of TLR2 and DC-SIGN in M2-like MDMs. (**D**) TLR2 and DC-SIGN expressions in MDDCs. (**E**) Secretion of TNF-α, IL-6, and IL-10 following day 6 LPS restimulation in monocytes treated with α-TLR2 or isotype control antibody prior to glycolipid stimulation (n=12). MFI = Mean Fluorescent Intensity, Stim. = Stimulation, Re-stim. = LPS restimulation on day 6. Seeding data is equivalent for all conditions, displayed in colors for comparison. Statistics: A-D: One-way Anova with Dunnett’s correction. E: paired t-test (Isotype control vs α-TLR2), *: p ≤ 0.05, **: p ≤ 0.01, ***: p ≤ 0.001, ****: p ≤ 0.0001

PIM_6_ and LAM have previously been reported to interact with TLR2 (Gilleron *et al*, 2006; Nigou *et al*, 2008). Thus, we introduced TLR2 blocking antibodies to monocytes prior to stimulations with PIM_6_, LAM, and combined TLR2 ligands Pam3CSK4 and Pam2CSK4 (Pam2/3CSK4). Monocytes were rested as before and TNF-α, IL-6, and IL-10 production was quantified after 24 hours of day 6 LPS restimulation. TLR2 blockade resulted in a significant, yet incomplete rescue of monocyte tolerization when compared to isotype control in cells stimulated with PIM_6_ and Pam2/3CsK4 (**Figure 2E).**

Blockade of Dectin-2, DC-SIGN, and Mincle resulted in no effect on PIM_6_-induced tolerization in monocytes **(Figure S2).**

### PIM_6_-Induced Tolerization is TLR2-Independent

Recent studies have shown that purified PIM_6_ may contain co-isolated lipopeptides (Belkai *et al*, 2026) , which are known to bind TLR2 (Fitzgerald & Kagan, 2020) and could be responsible for propagation of a tolerized response in myeloid cells. We therefore treated PIM_6_ and LAM with hydrogen peroxide, which is known to inactivate TLR2-lipopeptide interaction (Zähringer *et al*, 2008) while retaining PIM_6_ and LAM structures (Belkai *et al*, 2026). Hydrogen peroxide treatment eliminated TLR2 activity in PIM_6_ fractions and diminished activity from LAM fractions (**Figure 3A**), as recently reported (Belkai *et al*, 2026). No TLR4 activity was observed in treated or untreated PIM_6_ or LAM fractions, ruling out contaminating LPS as a source of the tolerized response (**Figure 3B**).

**Figure 3.**
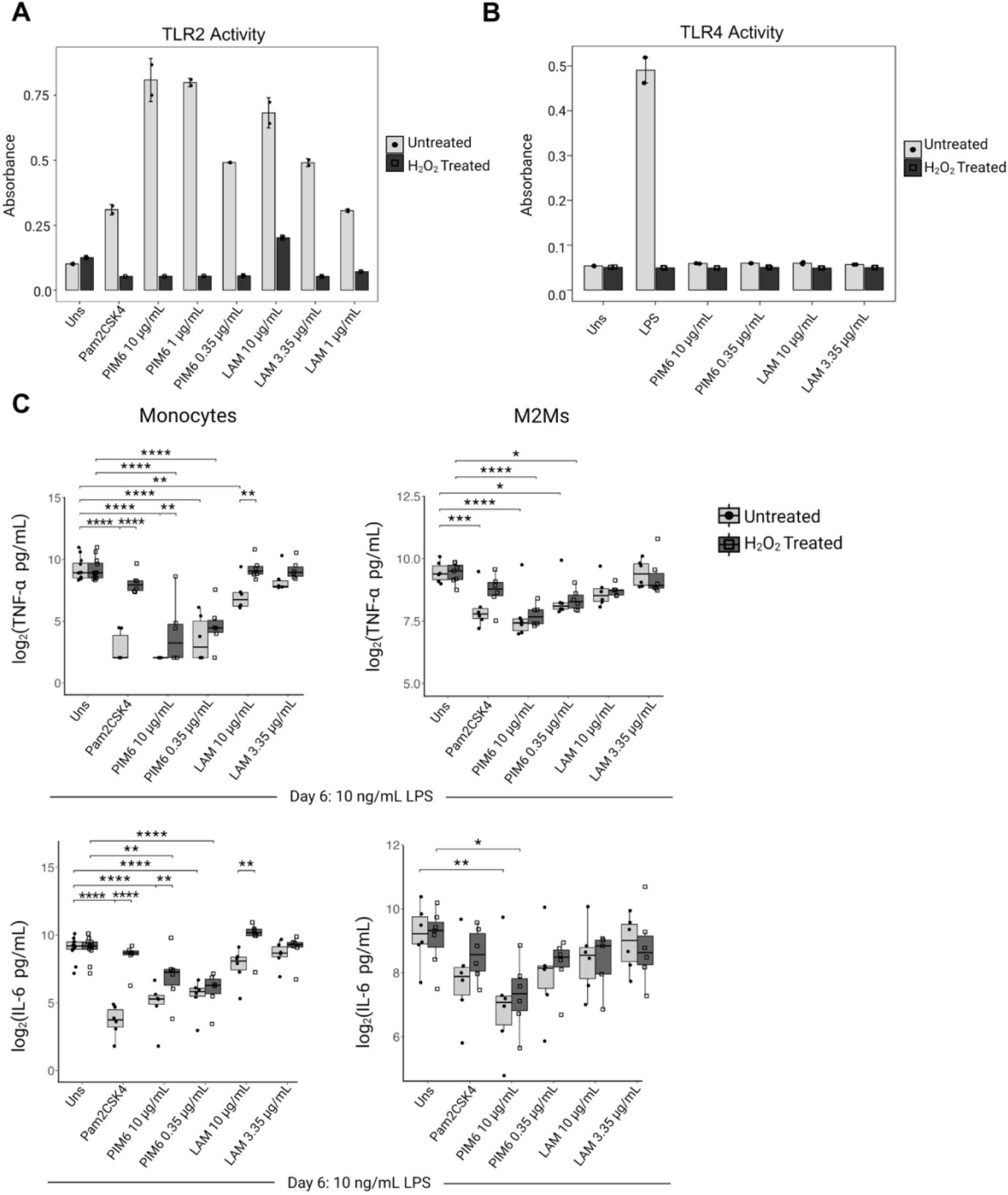
PIM_6_ Induces Tolerance in Myeloid Cells Despite Heterogenous Purified Fraction. (**A**) HEK TLR2 reporter cell lines were stimulated in two independent experiments with peroxide treatment and untreated lipid fraction. After 24 hours, TLR2 stimulation was assayed using a Quanti-Blu kit. (**B**) HEK TLR4 reporter cells line was stimulated in two independent experiments with peroxide treated or untreated lipid fractions. After 24 hours, TLR4 activation was assayed using a Quanti-Blu kit. (**C)** PIM_6_ fractions were treated with hydrogen peroxide to oxidize contaminating lipopeptides and were used in primary stimulation in monocytes and M2Ms. Cells were rested for 5 days and restimulated with LPS for 24 hours (n = 4-6). Stats: (**C**) Matched two-way ANOVA with Geisser-Greenhouse correction and Tukey’s multiple comparison test. Ns : p ≥ 0.05, *: p ≤ 0.05, **: p ≤ 0.01, ***: p ≤ 0.001, ****: p ≤ 0.0001.

To examine if lipopeptide-inactivated PIM_6_ could still induce immune tolerance in myeloid cells, we conducted the same stimulation protocol as in Figure 1 with hydrogen peroxide treated glycolipids and Pam2CSK4 to control for extinction of lipopeptide-TLR2 activity. Peroxide treatment of Pam2CSK4 almost completely eliminated the tolerized phenotype in monocytes and M2Ms (**Figure 3C**). Hydrogen peroxide treated PIM_6_ maintained its tolerizing effect, though a small difference in cytokine secretion from treated vs untreated PIM_6_ could be observed at high concentrations (**Figure 3C**). A small difference in TNF-α and IL-6 secretion from monocytes was also observed in high concentrations of treated vs untreated LAM, though neither condition resulted in a significantly dampened secretion when compared to unstimulated control. This is consistent with recent findings of retained TLR2 activity in high concentration of hydrogen peroxide treated LAM(Belkai *et al*, 2026). Collectively, these findings definitively identify PIM6 as a tolerizing ligand, irrespective of TLR2-lipopeptide interactions.

### PIM_6_ Transiently Enhances Bacterial Control in M.tb Infected M2Ms

To investigate if PIM_6_ or LAM-induced innate modulation could affect subsequent control of M.tb infection, M2Ms were pretreated with glycolipids for 24 hours and rested before infection at an MOI of 1 with H37Rv M.tb constitutively expressing GFP. M2Ms were used in the infection model to simulate the phenotype of naïve macrophages and representative images of day 6 post-infection cultures are exhibited in (**Figure 4A**). M2M cell viability and number were assessed immediately prior to infection to ensure comparable downstream comparison of ROS production, cytokine secretion, and CFU analysis (**Figure S4**).

**Figure 4.**
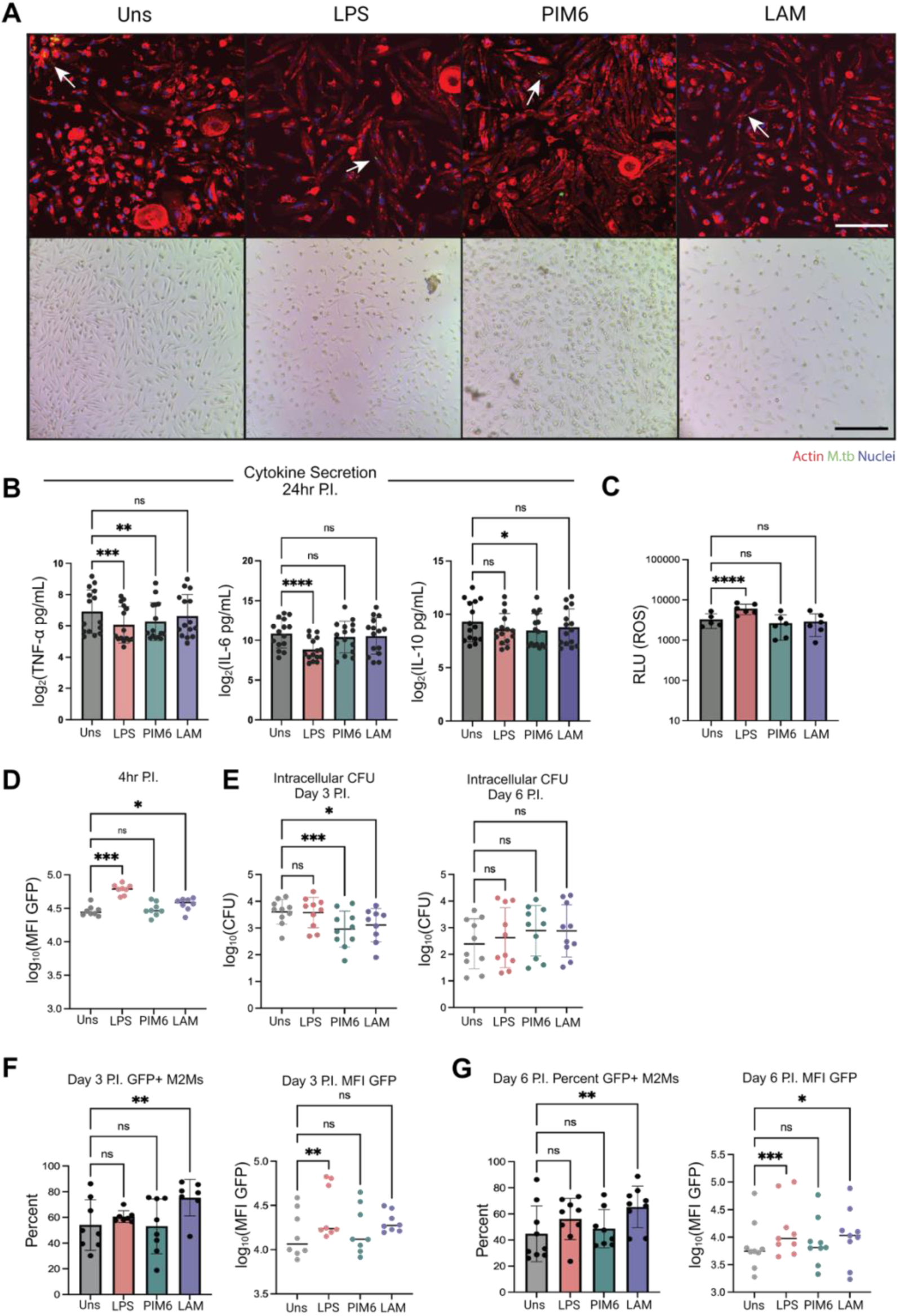
Effect of PIM_6_ and LAM Pretreatment on M.tb Infection Control. (**A**) Representative images of M2Ms 3 days P.I. with GFP-expressing H37Rv (MOI =1) after pretreatment with LPS, PIM_6_ or LAM. Arrows highlight intracellular M.tb. Top row: Fluorescent Images of infected M2Ms, stained with Phalloidin and DAPI. Scale bar represents 200 µm. Bottom row: brightfield images of infected M2Ms. Scale bar represents 400µm. (**B**) Cytokine secretion from M2Ms 24 hours after infection (n = 15). (**C**) ROS production 24 hours post infection (n = 6). (**D**) MFI of GFP M.tb in infected M2Ms immediately following infection (n =6). (**E**) Intracellular CFU recovered after 3 or 6 days after infection (n = 10). (**F & G**) Percentage of infected cells and bacterial burden via MFI GFP both 3 and 6 days after infection (n = 8). P.I = Post Infection. MOI = Multiplicity of Infection. CFU = Colony Forming Units. Statistics: Repeated measures one-way Anova or mixed e ffect model with Dunnett’s multiple comparison test. ns : p ≥ 0.05, *: p ≤ 0.05, **: p ≤ 0.01, ***: p ≤ 0.001, ****: p ≤ 0.0001.

Twenty-four hours following infection, culture supernatants were analyzed for TNF-α, IL-6, and IL-10 (**Figure 4B**). Consistent with the tolerized phenotype observed upon LPS restimulation, PIM_6_ pretreated conditions exhibited decreased concentrations of TNF-α and IL-10 compared to unstimulated control while LPS pretreated cells secreted less TNF-α, IL-6, and IL-10. Although ROS production was significantly upregulated in LPS treated conditions, the unaltered ROS production in PIM_6_ or LAM conditions indicated that subsequent bacterial control was independent of oxidative burst (**Figure 4C**). Immediately following infection, LPS and LAM pretreated M2Ms exhibited enhanced bacterial uptake (**Figure 4D**), consistent with previous works demonstrating that TLR signaling increases phagocytosis (Blander & Medzhitov, 2004).

On day 3 post infection, intracellular bacterial burden was significantly reduced in M2Ms pretreated with PIM_6_ compared with unstimulated controls (**Figure 4E**), demonstrating enhanced control of early bacterial growth. CFUs normalized to cell number per well yielded similar results (**Figure S4**). Flow cytometry of infected M2Ms on day 3 post infection showed no change in percentage of infected (GFP+) cells from PIM_6_ treatment while there was a significant increase in LAM treated cells (**Figure 4F**). The bacterial burden at day 3, measured by the mean fluorescent intensity of GFP was significantly increased in LPS treated cells, while PIM_6_ or LAM conditions showed no change (**Figure 4F**).

On day 6 post infection, intracellular CFU counts demonstrated no change in viable bacteria between treatment groups (**Figure 4E**). Flow cytometry analysis showed an increase in bacterial burden for LPS treated cells, and there was an increase in both percentage of infected cells and bacterial burden in the LAM treated group (**Figure 4F).** There was no change in extracellular CFUs between the conditions at day 3 or day 6 post infection (**Figure S4**). Together, these findings suggest that PIM_6_ pretreatment induces a transient reprogramming of macrophages that promotes early control of intracellular M.tb, although this effect was no longer evident by day 6 post infection.

## Discussion

In a previous study (Silva *et al*, 2021), we showed that myeloid cells in M.tb-immunoreactive (latently infected) individuals respond with a dampened cytokine production following stimulation with the M.tb glycolipids LAM and PIM_6_ compared to those who had uncontrolled infection in TB disease. This observation suggested that glycolipid-specific hypo-responsiveness in myeloid cells could be associated with infection control. We thus investigated whether mycobacterial glycolipids could induce a sustained tolerized phenotype consistent with innate immune memory (Divangahi *et al*, 2021; Saeed *et al*, 2014; Netea *et al*, 2020).

The present study is the first, to our knowledge, to incorporate mycobacterial glycolipids into a protocol to examine a memory-like modulation of the myeloid cell compartment, demonstrating that PIM_6_ plays a significant role in establishing a tolerized phenotype, most markedly in monocytes, and M2Ms. Innate immune memory can manifest as either trained immunity or innate immune tolerance, resulting in enhanced or reduced responses to secondary stimulation (Divangahi *et al*, 2021; Silva *et al*, 2021; Das *et al*, 2019).

While previous literature has demonstrated that exposure to M.tb glycolipids can downmodulate the myeloid immune response to inflammatory stimuli (Källenius *et al*, 2020; Correia-Neves *et al*, 2022; Nigou *et al*, 2001; Blanc *et al*, 2017; Silva *et al*, 2021; Garcia-Vilanova *et al*, 2019; Vergne *et al*, 2015; Bah *et al*, 2020), these studies administered M.tb glycolipids immediately before or simultaneously with immune challenge. With simultaneous stimulation and readout, it is unclear whether this downmodulation is a transiently impaired immune cell activation or a more long-lived response. In this study, we directly interrogate the capacity of mycobacterial glycolipids to induce tolerance in primary myeloid cells by resting cells for five days between glycolipid stimulation and LPS challenge.

We anticipated that decreased TNF-α and IL-6 production in PIM_6_ pretreated M2Ms would be especially susceptible to M.tb infection (Verrall, 2020). Instead, PIM_6_ pretreatment led to a significant decrease in culturable bacterial load during early infection. While flow cytometry analysis suggests that PIM_6_ did not affect bacterial uptake nor burden on day 3 and 6, CFU counts normalized for cell number further supported decreased culturable M.tb at day 3 post infection. This may be attributed to the upregulation of non-cytokine antimicrobial mechanisms not observed in our study, associated with, or unrelated to the observed tolerized cytokine secretion. Alternatively, macrophage activation state may attenuate the ability of M.tb to replicate (Liu *et al*, 2016), effectively uncoupling proinflammatory responses with antimicrobial function. While this protective effect was transient in our *in vitro* system, it could still be crucial in M.tb infection as the early innate response coordinates the antimicrobial activity of both the innate and adaptive immune systems.

LPS prestimulation also dampened cytokine secretion in macrophages upon M.tb infection, despite enhanced phagocytosis and Mtb uptake (Kirchhoff *et al*, 2025). While intracellular burden, assessed by GFP MFI, was increased at day 3 and 6 post infection, LPS CFU recovery was comparable to unstimulated control. This indicated that increased initial bacterial uptake was counterbalanced by enhanced intracellular control mechanisms, preventing culture of viable M.tb.

Despite LAM’s well documented effects on myeloid immunomodulation, and our own findings that PBMCs from individuals with TB infection are less reactive to LAM than those with TB disease, LAM did not enact a tolerizing effect in our cell types that we could detect with our protocol. Conversely to our PIM_6_ pretreated condition, LAM stimulated cells experienced higher percentage of infection at both timepoints and increased bacterial burden. We posit that, because LAM not only enhances bacterial uptake, but also integrates into the cell wall, that LAM could remain within the culture following the stimulation, thus contributing to virulent function of the bacteria (Goren *et al*, 1976; Kang *et al*, 2005; Sweet *et al*, 2010), and resulting in the enhanced intracellular bacterial persistence and replication.

Limitations of this study include the lack of donor IGRA status and the use of circulating monocytes. Though healthy blood donors recruited in the Stockholm region are likely to have limited prior exposure to M.tb, this was not formally assessed, and the potential contribution of previous exposure to experiment outcome cannot be excluded. Additionally, given that innate immune memory can persist long term and be established at the level of hematopoietic stem and progenitor cells, the tolerogenic effects of PIM_6_ observed in circulating myeloid cells may extend beyond the peripheral compartment, suggesting broader implications for immune functions.

Overall, these data present PIM_6_ as a complex tolerogenic immunomodulator with direct effects on M.tb infection in primary human macrophages. However, the exact mechanisms and receptors by which PIM_6_ induces a tolerized response to subsequent stimuli and infection remain unknown. In our study, functional blockade of DC-SIGN and Dectin-2, receptors previously reported to bind PIM6 (Driessen *et al*, 2009; Decout *et al*, 2018), did not alter the tolerizing effect of PIM6 in monocytes. Therefore, broader approaches, such as transcriptomic profiling by RNA-seq, are required to elucidate signaling mechanisms enacting this phenotype. Further studies should also assess whether lower-order molecules, such as PIM_2_ and PI, present as residual components within PIM_6_-enriched fractions, contribute to the observed tolerogenic effects.

## Materials and Methods

### Mycobacterial Glycolipid Isolation

The following reagents were obtained through BEI Resources, NIAID, NIH: *Mycobacterium tuberculosis*, Strain H37Rv, Purified Phosphatidylinositol Mannoside 6 (enriched fraction, hereafter referred to as PIM_6_), NR-14847 (Lot #: 70037069, 70054808), and *Mycobacterium tuberculosis*, Strain H37Rv, Purified Lipoarabinomannan (LAM), NR-14848 (Lot #: 70037070). Other PIM6 stocks were purified from H37Rv M.tb as previously described (Gilleron *et al*, 2006) .

### Cell Culture

Peripheral blood mononuclear cells (PBMCs) were isolated from freshly obtained buffy coats from Karolinska Institute Clinical Immunology and Transfusion Medicine and Stockholm Bloodbank via Ficoll Paque (Cytiva #17144002) density gradient centrifugation. 20 mL of buffy coat per donor was mixed 1:1 with room temperature dPBS (Sigma Aldrich #D8537) and carefully pipetted atop 15 mL Ficoll Paque Plus solution in a 50 mL Leucosep Centrifuge Tube (Greiner #227290), carefully making sure that the blood layered atop the Ficoll solution. Tubes were then spun for 15 minutes at 800 g with no brakes. The resulting buffy layer was pipetted into a separate 50 mL Falcon tube and washed with 50 mL room temperature dPBS twice, spinning down at 250 g for 10 min, half brake. After the final wash, the pellet was resuspended in 50mL dPBS and 10 µL of the suspension was mixed with 10 µL of Trypan Blue Stain (Gibco #15250061) for counting using a Countess II automated cell counter. The appropriate volume containing the number of PBMCs needed for monocyte isolation was spun down a final time at 300g for 10 min, full break.

Monocytes were isolated from PBMCs using a CD14 magnetic microbeads kit (Miltenyi #130-097-052), or isolated with RosetteSep Human Monocyte Enrichment cocktail (Stemcell #15028) or from plastic adherence as previously described (Nielsen *et al*, 2020; Domínguez-Andrés *et al*, 2021) . Purity of isolated monocytes was analyzed via flow cytometry, and a threshold of 95% purity was used for the following experiments. Purified monocytes were counted and seeded at a density of 1 x10^5^ cells/ well in a 96-well plate (Corning #3596) in complete RPMI made up from RPMI 1640 (Sigma-Aldrich #R0883) supplemented with 10% heat-inactivated fetal bovine serum (Gibco #A5256501), 2mM L-glutamine (Gibco #25030081), and 100 U/mL penicillin/streptomycin (Gibco #15140).

Monocytes were left resting overnight (∼16 hours) at 37°C and 5% CO_2_ before stimulation. M1Ms and M2Ms were differentiated in complete RPMI containing 50 ng/mL GM-CSF or M-CSF respectively and MDDCs were differentiated with 50 ng/mL GM-CSF and IL-4 over 6 days, changing media every 48 hours. Cells were then phenotypically characterized by flow cytometry through evaluation of CD200R, CD163, CD86, and CD64 (Mily *et al*, 2020) (**Table 2**).

LDH quantification to assay cell death was conducted according to manufacturer’s instructions (Promega, #J2380).

### Flow Cytometry

Cells were collected from suspension or detached from well plates using 4°C incubation with 5mM EDTA in PBS followed by scraping. Cells were washed twice in cold PBS prior to staining for 30 minutes with Live Dead Green (Invitrogen #L34969) or Aqua (Invitrogen #L34957) diluted 1:600 or 1:1200 in PBS respectively at 4°C. Cells were washed twice with PBS +2% FBS before administration of extracellular antibody cocktail with for 30 minutes at 4°C. Cells were washed again with PBS + 2% FBS three times followed by a 15 minute room temperature fixation with 4% Paraformaldehyde. For Infection experiments, fixation was extended to 12 hours to inactivate M.tb. Fixed cells were washed twice with PBS + 2% FBS and resuspended in 200 µL. Stained samples were analyzed on a Cytek Aurora 3 or 5 Laser full spectrum flow cytometer and samples were analyzed with FlowJo version 10.10.1.

### Stimulations

All cell types were stimulated with 10 ng/mL LPS (Invivogen #tlr1-b5lps), 350 ng/mL PIM_6_ (BEI #NR-14848), 25 µg/mL LAM (BEI NR14847), 25 µg/mL M.tb cell lysate (BEI #NR-14822), or were left unstimulated in complete RPMI for 24 hours. Wells were washed twice with warm dPBS to remove excess stimulant and topped with resting medium of complete RPMI. Cells were rested for 5 days, with a medium change on day 3. On Day 6, cells underwent secondary stimulation with 10 ng/mL of LPS for 24 hours. After 24 hours, supernatants were collected and frozen at –80°C for cytokine analysis.

### Functional Antibody Blocking

Monocytes were seeded as previously described and incubated with 5 μg/mL TLR2 blocking antibody (Invivogen #mab2-mtlr2) or a mouse IgG_1_ (ebiosciences #16-4714-85) isotype control for 4 hours at 37° C and 5% CO2. Cells were then stimulated with 350 ng/mL PIM_6_ or 5 ng/mL Pam2Csk4 (Invivogen #tlrl-pm2s-1) (TLR2/6 agonist) + 5 ng/mL Pam3Csk4 (Invivogen #tlrl-pms) (TLR2 1/6 agonist). After 24 hours of stimulation, the supernatant was frozen at -80° C for verification of TLR2 blockade with TLR2 agonists before cells were washed with warm dPBS and topped with resting medium (complete RPMI). Cells were cultured for 5 days, changing media on day 3 before restimulating with 10 ng/mL of LPS for 24 hours. Supernatants were then collected and stored at -80° C for cytokine analysis.

### Titration of PIM_6_ and LAM

Monocytes from four healthy donors were isolated and seeded at a density of 5×10^5^ cells/well into 24-well plates (Corning #CLS353226) and stimulated with 350, 140, 70, or 14ng/mL of PIM_6_ for 24 hours. Cells were then washed with warm dPBS and wells were topped with complete RPMI. Cells were rested for 5 days, changing media on day 3. On Day 6, cells were stimulated with 10 ng/mL of LPS for 24 hours. After 24 hours, cell culture supernatant was collected and stored at -80° C until cytokine analysis.

### Glycolipid Hydrogen Peroxide Reduction

Lyophilized PIM_6_, LAM, and Pam2CsK4 were reconstituted in 10% DMSO (Sigma #D4540) in MiliQ water and the desired amount was aliquoted into wells of a 96 well plate. Glycolipids were treated with 1% H2O2 (Fisher #H325) for 3 hours at 37°C in the well plate or MQ water for untreated controls. After treatment, the suspended glycolipid solutions were dried completely with forced air. Cells were seeded in appropriate wells.

### HEK Reporter Cell Line Experiments

HEK-Blue TLR2 (Invivogen #hkb-htlr2) and HEK-Blue TLR4 (Invivogen #hkb-htlr4) cell lines were maintained in DMEM (Sigma-Aldrich #D57962) supplemented with 2mM L-glutamine (Gibco #25030081), 10% heat-inactivated fetal bovine serum (Gibco #A5256501), 100 U/mL Pen-Strep (Gibco #15140), and 1X HEK-Blue Selection (Invivogen #hb-sel).

Each cell line was seeded at 50,000 cells per well in the previously prepared 96 well plate containing hydrogen peroxide treated glycolipids. Cells were incubated at 37°C and 5% CO_2_ for 24 hours before supernatant was collected and a Quanti-Blu Assay (Invivogen #rep-qbs) was carried out according to manufacturer’s instructions to detect TLR activity.

### Cytokine Quantification

Cytokine levels were quantified using the LEGENDplex™ Human Essential Immune Response Panel (Biolegend #740930) or the LEGENDplex™ HU Th22 Panel kit (Biolegend #741038). The assay protocol was followed as indicated by the manufacturer, and beads were run on a Cytek Aurora 3 Laser Spectral Flow cytometer. Spectral data was analyzed using the free LEGENDplex ™ Qognit Software Analysis Suite.

### M.tb Culture

An H37Rv strain of M.tb constitutively expressing GFP under the smyc reporter was provided by the Karolinska Institutet Bomedicum BSL3 facility. GFP M.tb was cultured in Middlebrook 7H9 broth (BD #271310) containing 10% ADC enrichment supplement (BD #212352) 0.2% Tween 80 (Fisher #9005-65-6), 0.2% glycerol (Thermo Fisher Scientific #BP229-1) and 20 μg/mL Kanamycin (Thermo Fisher Scientific #J60668.03). Broth cultures were shaken at 150 rpm and incubated at 37°C with 5% CO_2_. Passages P3-5 were used for infection experiments and optical density at 600 nm was used to calculate culture concentration for subsequent infections.

### M.tb Infection of MDMs

Primary monocytes were isolated and seeded at 10.5x 10^4^ cells/cm^2^ into either a Falcon 8 Chamber culture slides (Falcon #354108), or a 24 well plate (Corning #CLS353226) and differentiated into M2Ms as previously described. Cells received a primary stimulation of 350 ng/mL PIM_6_, 3.35 ug/mL LAM, or 10ng/mL LPS for 24 hours before stimulus was washed and cells were rested for 5 days. Cells were infected with a strain of H37Rv M.tb constitutively expressing GFP (MOI=1) for 4 hours before washing and topping with complete RPMI.

ROS production was measured using the ROS-Glo H_2_O_2_ Assay (Promega G8820) according to manufacturer’s instructions at 24 hours following wash of extracellular M.tb.

24 hours after infection, supernatant was collected and analyzed for cytokine secretion. Days 3 and 6 post infection, both supernatant and cell lysate were plated for extracellular and intracellular CFUs count respectively on Middlebrook 7H11 (BD #212203) agar supplemented OADC (BD # 212351). Plates were incubated for 21 days at 37°C before quantification. CFU values were calculated from dilutions containing 10-100 colonies per plate.

Days 3 and 6 post infection, cells were also lifted with cold PBS containing 5 mM EDTA. Macrophages were stained for flow cytometry (**Table T1).**

### Fluorescent Imaging

On days 3 and 6 post infection, cells were fixed and M.tb was neutralized overnight with 4% PFA at 4° C (Thermo Fisher Scientific #J19943.K2). Cells were washed thrice with PBS and permeabilized 15 min with 0.1% Triton X-100 (Invitrogen #HFH10) in PBS at room temperature. Cells were again washed with PBS and stained for actin (Phalloidin Rhodamine, (Invitrogen #R415) 1:400) and DAPI ( (Invitrogen #62248) 1:10,000 in PBS). Cells were washed thrice with PBS, once with DI water, and a cover glass was mounted using Prolong Gold antifade mounting medium (Invitrogen #P10144). Fluorescent imaging of infected M2Ms was conducted with a Zeiss LSM 800 laser scanning confocal microscope using a 20X objective. Tile stitching was conducting on Zen Blue v2.3 software. Further channel merging was conducted using Image J (Fiji).

## Data Availability Statement

This study includes no data deposited in external repositories.

## Author Contributions

M.S.G., C.S., G.K., and J.N designed the study. M.S.G., L.K., and C.S.S. performed experiments. Creation of figures and tables was conducted by M.S.G.. Data analysis was conducted by M.S.G., G.K., C.S., and M.C.N.. J.N. and M.G. provided key resources. M.S.G., G.K., M.C.N., and C.S. wrote the manuscript.

All authors contributed to the revision and approved the submitted version.

## Acknowledgments

We would like to thank Jerome Nigou for generously supplying the HEK-Blue reporter cell lines used in this manuscript. We would also like to thank Martine Gilleron for supplying us with purified PIM_6_ fractions. We further thank the lab of Martin Rottenberg for providing GFP expressing H37Rv *Mycobacterium tuberculosis* for this manuscript.

This study was supported by the Swedish Heart and Lung Foundation (HLF) (2022-0181 to G.K.), the Swedish Research Council (VR) (2020-03602 to G.K.), and Karolinska Institutet KID funding (2022-00880 to C.S.).

We further thank BEI Resources NIAID, NIH for supplying purified glycolipids (PIM6: NR-14848, LAM: NR-14847).

We thank the Karolinka Institutet Center for Molecular Medicine Flow Cytometer Core Facility for access to flow cytometers and the Karolinska Institutet Biomedicum Imaging Core for access to confocal microscopes.

The figures in this manuscript were created using Biorender.

Figure 1: Created in BioRender. Gower, M. (2027) https://BioRender.com/nm3zbk5.

Figure 2: Created in BioRender. Gower, M. (2027) https://BioRender.com/t8h8jj1.

Figure 3: Created in BioRender. Gower, M. (2027) https://BioRender.com/aryey5l.

Figure 4: Created in BioRender. Gower, M. (2027) https://BioRender.com/7l1huys.

