## Supplementary Information for "PIM_6_ Induces Tolerance and Confers Protection against *Mycobacterium tuberculosis* in Primary Macrophages"

### Supplemental Information

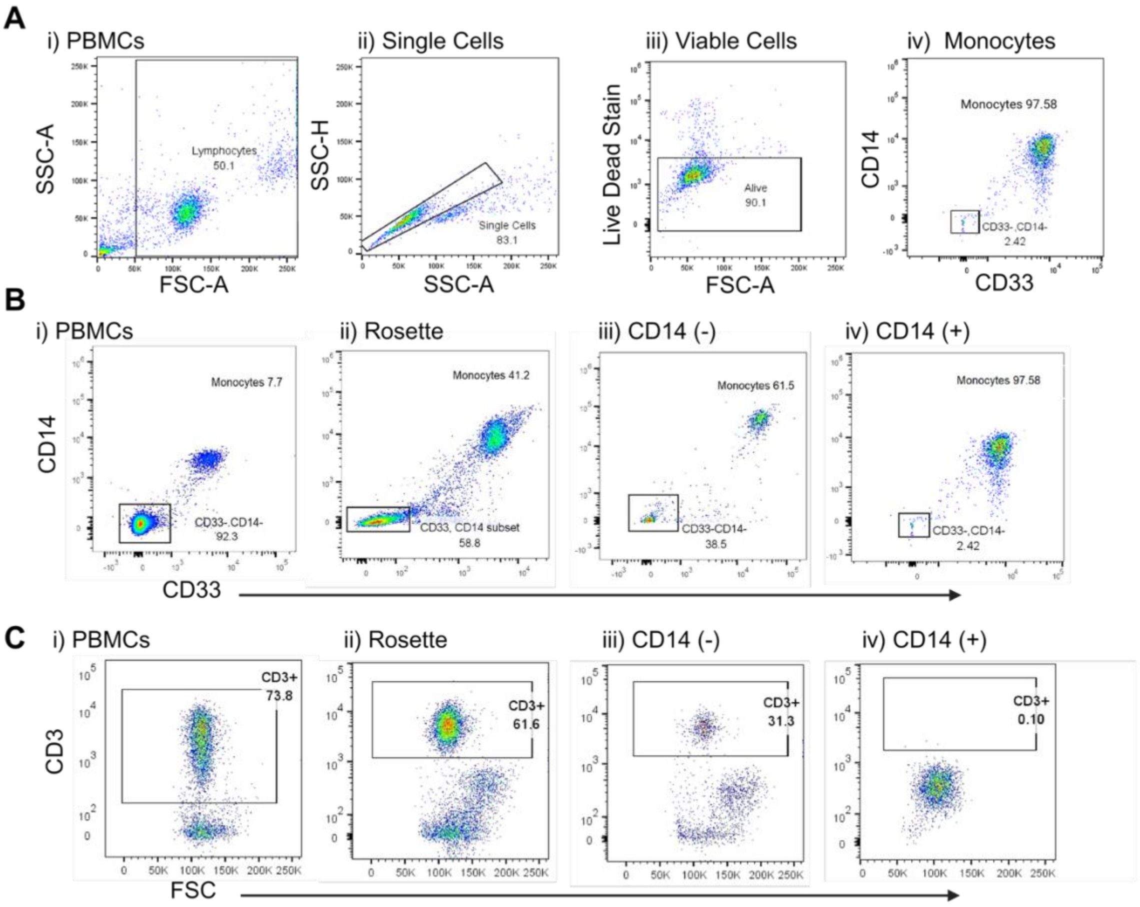


**Figure S1. Validation of T cell depletion in CD14+ magnetic bead isolation and support for maintained cell viability throughout assay.** (**A**) Gating strategy for evaluation of monocyte isolation purity. (**B**) Evaluation of monocyte purity from different methods of CD14+ monocyte isolation via Rosette Sep negative selection, CD14 negative magnetic isolation, and CD14 positive isolation. (**C**) Evaluation of CD3+ cell populations in PBMCs and differing isolation methods.

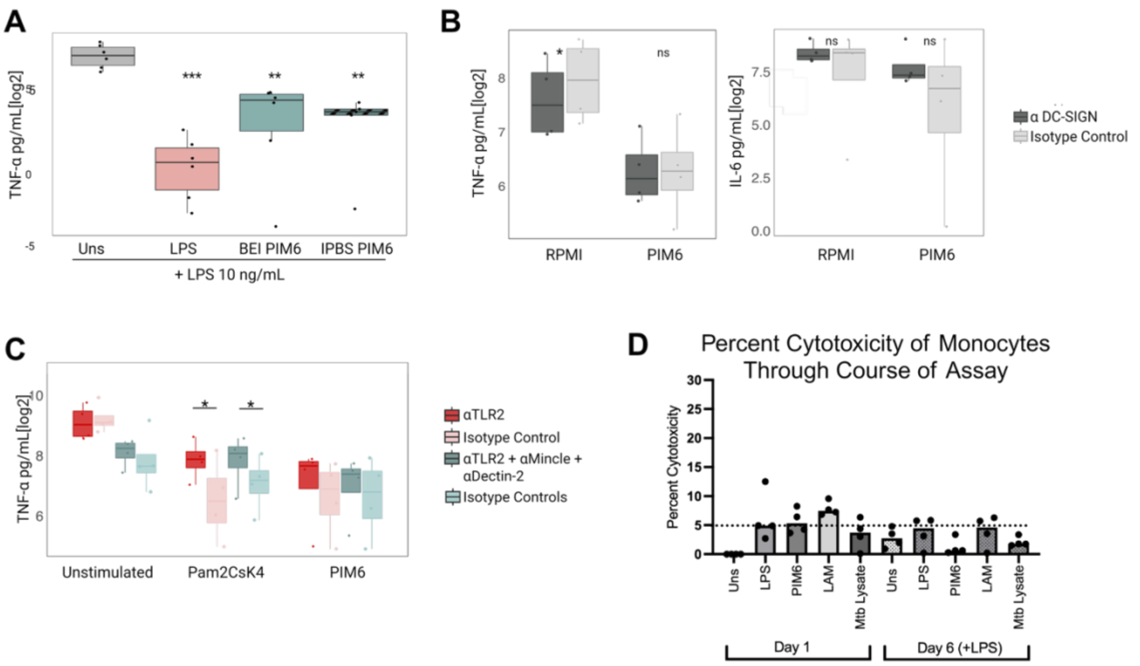


**Figure S2. PIM6 isolated from various sources indues tolerance in monocytes independent of DC-SIGN, Mincle, and Dectin-2 signalling.** (**A**) Monocytes were left in RPMI or stimulated with 10 ng/mL LPS, 350 ng/mL BEI PIM6, or 350 ng/mL PIM6 isolated at Universite de Toulouse. Cells were subsequently rested for 5 days before administering 10 ng/mL LPS on day 6. TNF-α secretion in the supernatant was analyzed (n = 6). (**B**) Monocytes were pretreated with a DC-SIGN blocking antibody prior to PIM6 stimulation. TNF-α and IL-6 were quantified following LPS (n = 4). (**C**) Monocytes were stimulated with Pam2CsK4 or PIM6 in the presence of α-TLR2 or combined α-TLR2+ α-Mincle + α-Dectin-2. TNF- α was quantified in the supernatant day 6 post LPS restimulation (n=4). (**D**) Supernatants were collected following 24-hour primary stimulation on day 1, and day 6 LPS stimulus from all stimulant conditions. Cell Cytotoxicity was calculated following lactate dehydrogenase quantification as the (($LDH_{Stimulant}$ – $LDH_{RPMI}$ )/ $LDH_{RPMI}$ x 100). Horizontal bar at 5% cell cytotoxicity for visualization (n = 4). Statistics: (A) Paired one-way anova with Dunnett's correction, unstimulated as reference group. (B & C) Pairwise comparisons were conducted using a t-test with Bonferroni correction. $p > 0.05$, *: $p \leq 0.05$, **: $p \leq 0.01$, ***: $p \leq 0.001$.

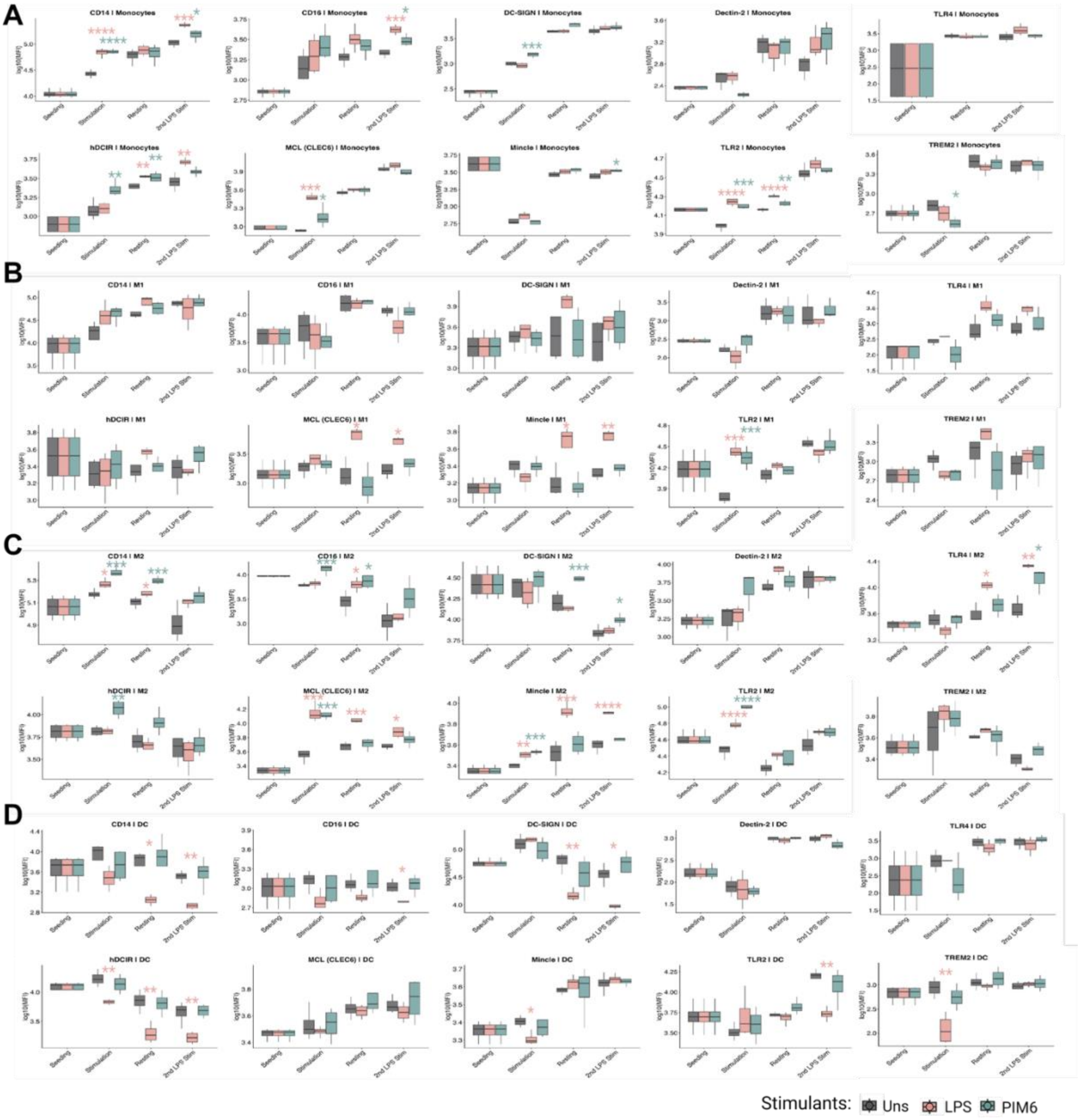


**Figure S3. Pattern recognition receptors in cells treated with LPS and PIM6.** Mean fluorescent intensity of CD14, CD16, DC-SCIGN, Dectin-2, hDCIR, MCL, Mincle, TLR2, TLR4 and TREM2 at seeding, after primary stimulation with LPS or PIM6, after 5 days of rest, and after LPS restimulation in (**A**) Monocytes, (**B**) M1Ms, (**C**) M2Ms, and (**D**) MDDCs. Statistics: paired t-test, *: p ≤ 0.05, **: p ≤ 0.01, ***: p ≤ 0.001, ****: p≤ 0.0001(n = 4).

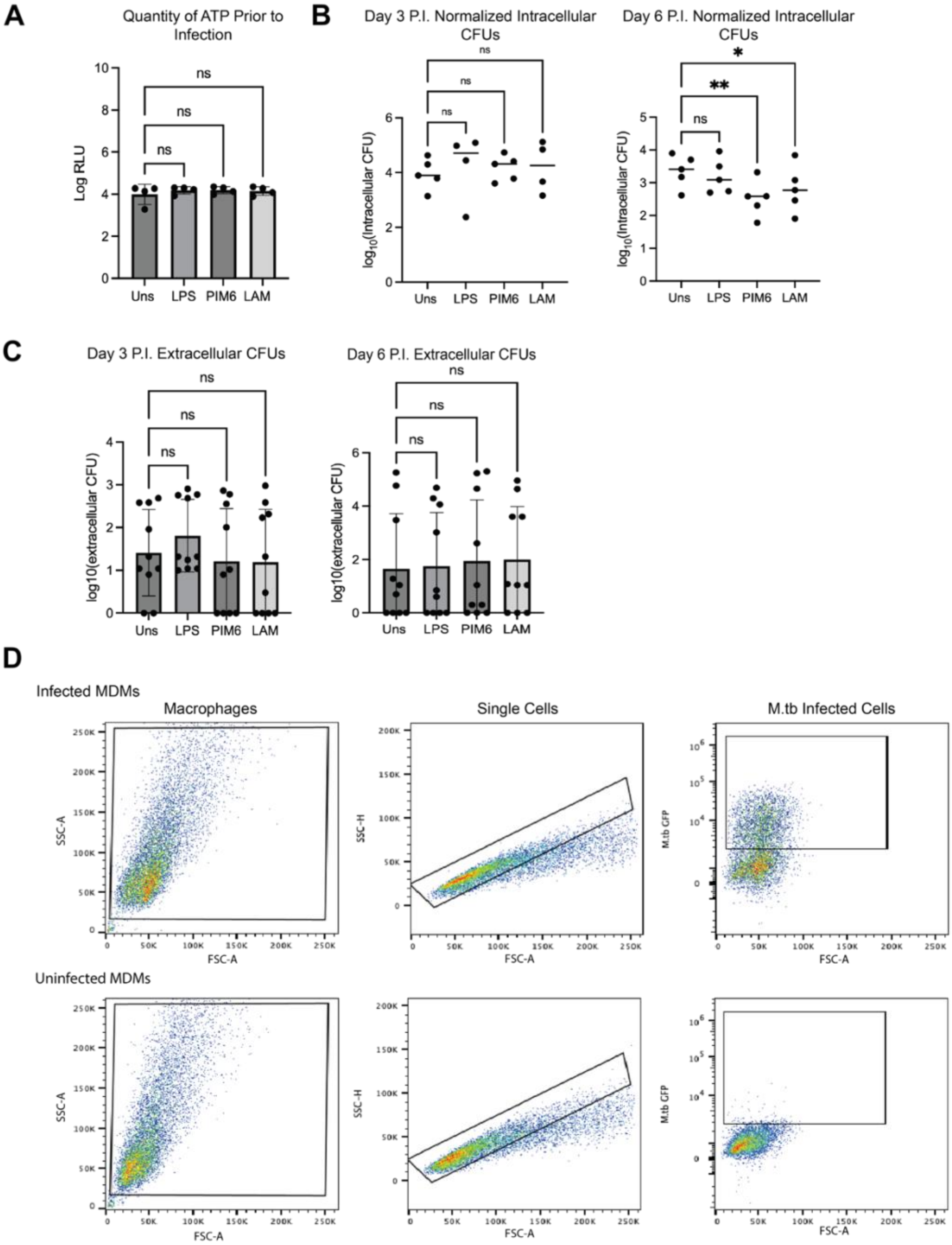


**Figure S4. M2M viability is equivalent across treatment groups after primary stimulation, immediately prior to M.tb infection, extracellular CFU quantification, gating strategy for percentage of infected M2Ms.** (**A**) ATP quantity in M2M cultures we assessed using Promega Cell Titer Glo kit to control for number of cells and cell viability (n=4 donors). (**B**) Counted CFUs were normalized against cell number per well counted by the ImageJ plugin Cell Counter from 20X brightfield images. (**C**) Extracellular CFU counts from day 3 and 6 post infection in infected M2Ms (n=10). (**D**) Representative gating strategy of infected or uninfected M2Ms at day 6 post infection. Statistics: Repeated measures one way ANOVA or mixed effects model with Dunnett correction.

**Table S1. Flow Cytometry Panel for CD14+ Monocyte Purity**

| Target/ Fluorophore | Company | Clone | Dilution |
|---|---|---|---|
| CD33 BV421 | BD Horizon | WM53 | 1:320 |
| CD14 PE | BD Pharmingen | MφP9 | 1:200 |
| CD163 BV650 | BD Horizon | GHI/6I | 1:40 |
| HLA-DR BV605 | BD Horizon | G46-6 | 1:120 |
| CD200R PE-Cy7 | Biolegend | OX-108 | 1:80 |
| CD64 APC Fire 750 | Biolegend | 10.1 | 1:40 |
| CD86 PE-Cy5 | Biolegend | IT2.2 | 1:90 |
| Live Dead Fixable Green Dead Cell Stain | Thermo Fisher Scientific | - | 1:1,200 |

**Table S2. Antibodies for pattern recognition receptor staining (multiple separate panels).**

| Target/ Fluorophore | Company | Clone | Dilution |
|---|---|---|---|
| TREM2 APC | R&D Systems | 237920 | 1:100 |
| DC-SIGN BB515 | BD Horizon | DCN46 | 1:120 |
| CD14 PE-Cy7 | BD Pharmigen | MφP9 | 1:300 |
| CD16 AF700 | Biolegend | 3GB | 1:120 |
| CLEC6 PE | Biolegend | 9B9 | 1:300 |
| Mincle AF488 | R&D Systems | 2455C | 1:100 |
| hDCIR AF647 | R&D Systems | I3-612 | 1:40 |
| TLR2 PE | BD Pharmigen | 11G7 | 1:100 |
| TLR4 BV421 | BD Horizon | TF901 | 1:120 |
| Dectin-2 AF647 | R&D Systems | 545925 | 1:100 |

**Table S3 Antibody Staining Panel for flow cytometry analysis of Infected Macrophages.**

| Target/ Fluorophore | Company | Clone | Dilution |
|---|---|---|---|
| CD33 BV421 | BD Horizon | WM53 | 1:320 |
| CD163 BV650 | Biolegend | GHI/6I | 1:40 |
| CD64 BV785 | Biolegend | 10.1 | 1:100 |
| TLR2 PE | BD Pharmigen | 11G7 | 1:100 |
| CD14 PEcf594 | BD Horizon | 63D3.r Mab | 1:120 |
| CD200R PE-Cy7 | Biolegend | OX-108 | 1:80 |
| DC-SIGN APC | BD Pharmigen | DCN46 | 1:120 |
| Live Dead Fixable Dead Cell Stain Aqua | Invivogen | - | 1:600 |
